# Expansion of modern humans over the world: The origin when considering non-linearity

**DOI:** 10.1101/2023.03.11.532168

**Authors:** Zarus Cenac

## Abstract

Types of diversity have been known to decline linearly with the rise of geographical distance from Africa. Declines have helped to suggest the area in Africa which holds the origin for the global expansion of modern humans. Research has, at times, explored if there is a non-linear relationship between diversity and distance from Africa. A previous suggestion was that non-linearity could affect where the expansion appears to have originated. Linear analysis with Y-chromosomal microsatellite heterozygosity has been contrary to the expansion from Africa, instead indicating an origin involving Asia; could this be attributable to non-linearity? The present study looked into whether there are non-linear relationships between distance and diversities, and approximated where the expansion began. This study used diversities from previous research – genetic (autosomal, X-chromosomal, Y-chromosomal, and mitochondrial) and cranial shape. The Bayesian information criterion was the statistic for comparing linear and non-linear (quadratic) models to indicate if there is a non-linear relationship. This criterion was also used to estimate where the expansion launched from. Autosomal microsatellite heterozygosity favoured a non-linear relationship. This may be due to South American populations. Mitochondrial diversity suggested non-linearity too, but not when minimum temperature was controlled for. Whilst non-linear relationships indicated that the expansion had its start in Africa, for autosomal microsatellite heterozygosity, the area of origin appeared to be rather affected by the type of model (linear or non-linear). Other diversities (e.g., Y-chromosomal) supported linear relationships. Therefore, non-linearity does not seem to explain Y-chromosomal microsatellite heterozygosity being unexpressive of the global expansion.

## Introduction

There has been discussion regarding how modern humans populated the world (e.g., Betti et al., 2009; Ramachandran et al., 2005). Africa has backing for being where modern humans emerged (Stringer, 2014) and globally expanded from (Ramachandran et al., 2005). Support for Africa being the origin of the expansion has involved certain types of diversity (e.g., Manica et al., 2007). A link between expansion and diversity can be understood through bottlenecks – conceivably, diversity would be lowered via bottlenecks, with more bottlenecks having been met as the expansion extended further away from its origin (Ramachandran et al., 2005). As distance from Africa amasses, sorts of diversity do lessen (e.g., Balloux et al., 2009; Betti et al., 2009, 2012, 2013; Betti & Manica, 2018; von Cramon-Taubadel & Lycett, 2008). Linear declines appear evident in genetic elements (Balloux et al., 2009) such as the heterozygosity of autosomal microsatellites (Prugnolle et al., 2005) and autosomal single nucleotide polymorphism (SNP) haplotypes, X- and Y-chromosomal microsatellite heterozygosities, and mitochondrial diversity (Balloux et al., 2009). Also, linear decreases look present in diversity within the skeleton (Betti et al., 2009, 2012; von Cramon-Taubadel & Lycett, 2008) such as in cranial form (Betti et al., 2009) and cranial shape (von Cramon-Taubadel & Lycett, 2008).

## Origin area

Attention has been given to where in Africa the expansion has its origin (e.g., Ray et al., 2005; Tishkoff et al., 2009). Consensus has not been reached on which region(s) of the continent the expansion originated from (Cenac, 2022b). Estimation of this origin has happened through different methods (e.g., Manica et al., 2007; Ramachandran et al., 2005; Ray et al., 2005; Tishkoff et al., 2009). For example, the area inside of which the origin resides may be estimated as i) the location which from which distance best predicts (the decline of) diversity, *and* ii) other locations from which distance is comparably good at predicting diversity (e.g., Betti et al., 2009; Manica et al., 2007; see also Cenac, 2022b) – see Figure 1A. The area can be referred to as the expansion origin area, and the location from which the decline appears strongest can be called the peak point (Cenac, 2022b). The peak point can be found using a measure called the Bayesian information criterion (BIC) – the lower the BIC, the better distance is at predicting diversity (Manica et al., 2007). As for locations that are equivalently effective at predicting diversity as the peak point is, BICs can be used for that too – locations giving BICs within four BICs of the peak point are (alongside the peak point) included in the origin area, unlike other locations (Manica et al., 2007) (see Figure 1A for instance). As an example, in Figure 1A, i) diversity is predicted *best* when distance to populations is from the peak point, ii) distance from another location inside the expansion origin area (i.e., within the dark grey area) to populations is similarly as good at predicting diversity as distance from the peak point, and iii) distance from locations outside the origin area (lighter grey) are not as good.

**Figure 1.**
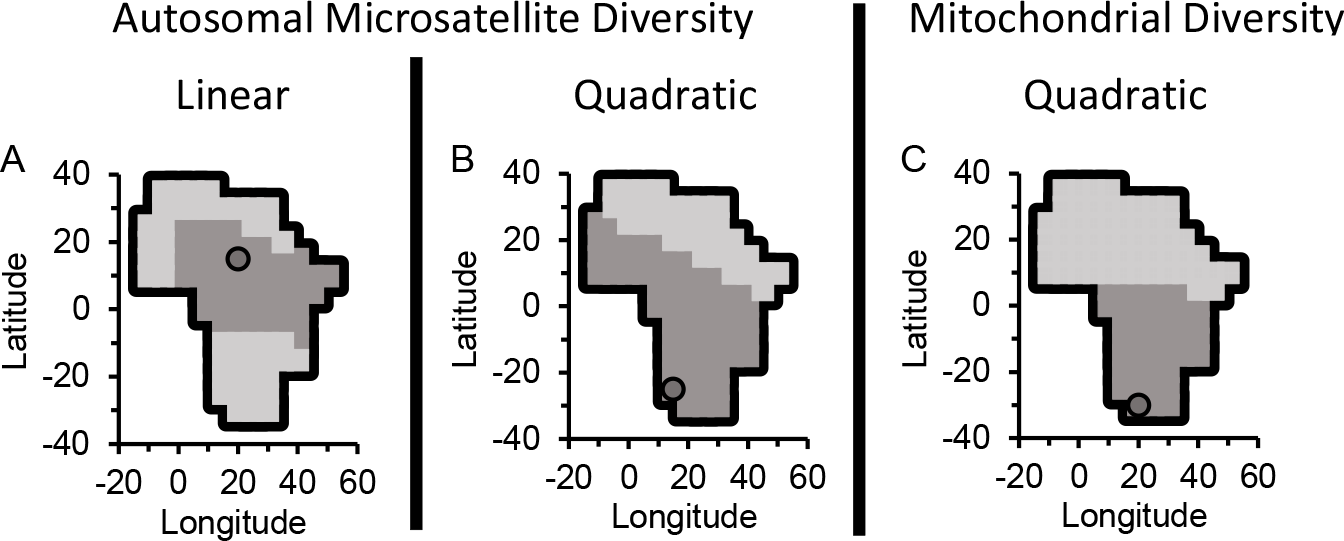
Origin Areas When Using BICs. *Note*. The origin area (darker grey) is shown, part of which is the peak point (circle). Regarding autosomal microsatellite heterozygosity of 50 HGDP-CEPH populations (Pemberton et al., 2013), the origin area using linear models (Figure 1A) appears to differ from the area estimated with quadratic models (Figure 1B). Using autosomal microsatellite heterozygosity in linear models, prior research has presented the origin area (Manica et al., 2007) and peak point (Ramachandran et al., 2005). Those estimates appear different than the ones in 1A. Utilising HGDP-CEPH data, those previous studies used 54 (Manica et al., 2007) and 53 population datapoints (Ramachandran et al., 2005), which are different numbers of datapoints compared to the present study, so some variation in estimates is to be expected. The origin area for mitochondrial diversity (quadratic) was estimated using mitochondrial diversity presented in Balloux et al. (2009). Although lands outside of mainland Africa are not shown in Figure 1, it was tested to some extent if they are part of the origin area of the expansion (see *Method*) – they were not found to be.

## Linearity

Using linear models, the origin area is wholly inside of Africa regarding autosomal microsatellite heterozygosity (Manica et al., 2007). A preprint using linear models found that Africa holds the origin area concerning autosomal SNP haplotype heterozygosity, X-chromosomal microsatellite heterozygosity, mitochondrial diversity, and cranial shape diversity, however, Africa was not in the origin area at all when it came to Y-chromosomal microsatellite heterozygosity (the area could very well be in Asia exclusively) (Cenac, 2022b).

The use of linear models when non-linear models (e.g., non-monotonic, like quadratic) are more appropriate may impact the estimation of where the expansion started from (Liu et al., 2006; see also Cenac, 2022b). Linearity has been considered in a number of studies, whether using simulations (DeGiorgio et al., 2009; Deshpande et al., 2009; Liu et al., 2006) or biological diversity (e.g., Betti et al., 2009). Regarding cranial form diversity, a non-linear trend is not supported for female crania (Betti et al., 2009). For male crania, support for non-linearity was found when counting outliers, although not when outliers were discounted (Betti et al., 2009). For sub-Saharan African populations, a previous study (a preprint) favoured there being a non-linear relationship (quadratic to be more specific) between distance from southern Africa and autosomal microsatellite heterozygosity (Cenac, 2022b). Indeed, autosomal microsatellite heterozygosity appeared to decrease only when distance from the origin reached about 2,000-3,000 km (and the diversity may be increasing between the origin and 2,000-3,000 km) (Cenac,

2022b). The non-linearity has different potential explanations, including admixture (between modern humans) happening particularly early in the expansion, thereby opposing the decline in heterozygosity which would arise from population bottlenecks (Cenac, 2022b). Nonetheless, whether a linear model or a quadratic model was used, southern Africa gained more support for being the origin compared to other regions of the continent (Cenac, 2022b). As for mitochondrial diversity, this diversity has a linear fall as distance from Africa broadens (Balloux et al., 2009). This has been found with hypervariable segment I and with the whole genome (absent of hypervariable segment I as well as II) (Balloux et al., 2009).

Estimation of the origin area has occurred regarding the whole genome (with the subtraction of those hypervariable segments), and this origin area is solely within Africa (Cenac, 2022b). Nevertheless, going from Figure 2C of Balloux et al. (2009), which presented distance from Africa against whole genome (minus both segments) mitochondrial diversity, it seems that there may be a quadratic relationship between distance from Africa and mitochondrial diversity. Additionally, mitochondrial diversity is related to climate – this sort of diversity increases linearly with minimum temperature (Balloux et al., 2009), so it could be that the potential non-linearity (regarding mitochondrial diversity and distance) may actually reflect the relationship between mitochondrial diversity and climate instead of the expansion.

**Figure 2.**
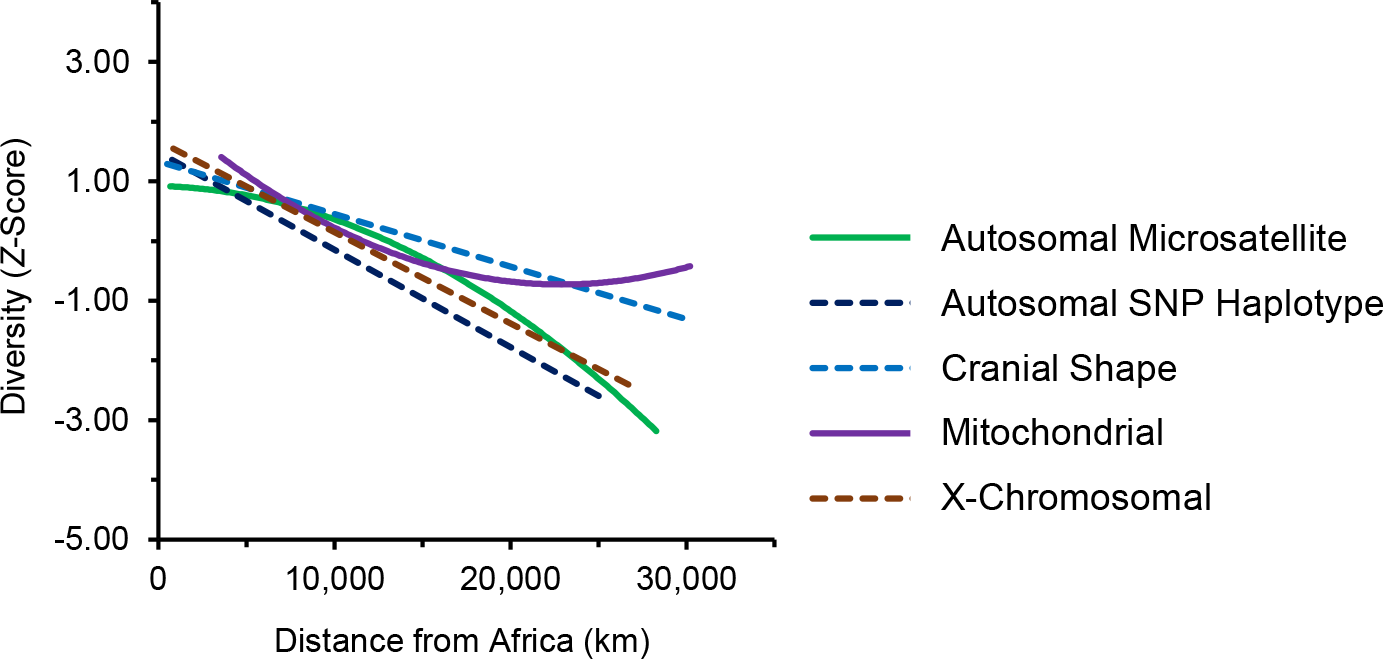
Trendlines for Diversities. *Note*. For Figure 2, diversities were *z*-score-standardised (within each type of diversity) so that trendlines regarding distance and diversity could be placed onto a common graph. Distances (Cenac, 2022b) were from peak points found in the present study or Cenac (2022b) for different types of diversity (for autosomal SNP haplotype heterozygosity, San were included).

Given preceding studies (Balloux et al., 2009; Betti et al., 2009; Cenac, 2022b; DeGiorgio et al., 2009; Deshpande et al., 2009; Liu et al., 2006), the present study examined if a quadratic trend is supported for different diversities: autosomal heterozygosity (microsatellite and SNP haplotype), allosomal heterozygosity (X- and Y-chromosomal), mitochondrial diversity, and cranial shape diversity. Because mitochondrial diversity is associated with a measure of minimum temperature (Balloux et al., 2009), there could be a point in finding what the peak point and origin area are when mitochondrial diversity is adjusted for minimum temperature (Cenac, 2022b); this was done in the present study, in which it was tested if an adjustment for minimum temperature is worthwhile amongst linear and quadratic models. Additionally, given the association between mitochondrial diversity and minimum temperature (Balloux et al., 2009), it was seen if non-linearity is indicated in mitochondrial diversity when this diversity is adjusted for minimum temperature. Y-chromosomal microsatellite heterozygosity does not point to an African origin of expansion when using linear models, and instead may suggest Asia, and so Y-chromosomal microsatellite heterozygosity is not a marker of the expansion (from Africa) (Cenac, 2022b). However, if non-linearity affects estimation of the origin (Liu et al., 2006), it could be worth seeing if non-linearity is found with Y-chromosomal diversity; the present study observed whether a non-linear trend appears.

## Method

R Version 4.0.5 (R Core Team, 2021) was utilised in the current study for calculations as was Microsoft Excel. BICs were calculated in Microsoft Excel through the use of a formula shown in Masson (2011) and by using R Version 4.0.5. Bartholomew illustrated reference atlas of the world (1985) was used to see which country the peak points (of quadratic models) are in and in which country/countries the origin areas are located. Southern Africa was regarded as the region described in Choudhury et al. (2021) when referring to origins.

## Genetic

Balloux et al. (2009) presented autosomal SNP haplotype heterozygosity (from Li et al., 2008, who calculated them from the HGDP-CEPH), both X- and Y-chromosomal microsatellite heterozygosities determined from the HGDP-CEPH, and mitochondrial diversity calculated from the mtDB alongside GenBank (whole genome with hypervariable segment subtractions); the present study used these diversities, each having 51 populations worldwide, presented in Balloux et al. (2009). To echo Cenac (2022b), with respect to mitochondrial diversity – Balloux et al. (2009) do cite Ingman and Gyllensten (2006) pertaining to the mtDB, and one can refer to Benson et al. (2013) regarding GenBank and Ingman and Gyllensten (2006) concerning the mtDB. Autosomal microsatellite heterozygosity (50 populations across the globe) was used from Pemberton et al. (2013); Pemberton et al. calculated the diversities from HGDP-CEPH data. Previous research (Cenac, 2022b; Pemberton et al., 2013; Tishkoff et al., 2009) was consulted in order to use populations which were not featured in autosomal microsatellite analyses (i.e., sub-Saharan African populations) in Cenac (2022b); this global analysis with autosomal microsatellite heterozygosity in the present study was used to see how the trend is worldwide, with different data being used than Cenac (2022b) for generalisability. Autosomal microsatellite diversity from HGDP-CEPH data is in a table in Balloux et al. (2009) to two decimal places. Instead, the present study used diversity from a table in Pemberton et al. (2013) because Pemberton et al. presented the diversity to a greater number of decimal places (three decimal places). Concerning mitochondrial diversity, minimum temperature measurements for populations are in Balloux et al. (2009), and the present study employed these measurements.

## Cranial

Regarding cranial shape diversity, von Cramon-Taubadel and Lycett (2008) and Cenac (2022b) used the Howells (1973, 1989, 1995, 1996) dataset (the Howells dataset is at http://web.utk.edu/~auerbach/HOWL.htm); the present study used the cranial shape diversity (of 28 populations) calculated in Cenac (2022b) as more decimal places are available than in von Cramon-Taubadel and Lycett (2008) Table 1. Whilst the Howells data has males in 28 populations, females do not feature in each of the 28 (von Cramon-Taubadel & Lycett, 2008), so population diversity of males was used rather than the diversity of females. Cranial shape diversity may represent the expansion more adeptly than cranial form diversity does (Cenac, 2022b), so cranial shape diversity was utilised instead of cranial form diversity in the current study.

**Table 1.** Number of Populations at Shorter Distances From the Peak Point.

| Diversity | < 2,000 km | < 3,000 km |
| --- | --- | --- |
| Autosomal microsatellite | 1 | 1 |
| Autosomal SNP haplotype | 4 | 4 |
| Cranial shape | 2 | 2 |
| Mitochondrial | 0 | 0 |
| X-chromosomal | 1 | 5 |

*Note.* As in Figure 2, distance was from the peak point for the relevant type of diversity, with peak points being determined in Cenac (2022b) or the current study. Given findings with diversity in sub-Saharan Africa (Cenac, 2022b), the particular distance categorisations of < 2,000 km and < 3,000 km in Table 1 were chosen.

## Distances and figures

Geographical distances in Cenac (2022a, 2022b) were made use of. Axes labels in figures follow Cenac (2022b) where appropriate. Peak point symbols are as in Cenac (2022b). The Africa representation in Figure 1A, 1B, and 1C is adapted from Cenac (2022b) (Cenac, 2022b, used locations presented in Figure 4 of Betti et al., 2013, and so the representation has its origins in Betti et al., 2013). Previously, the cranial diversity within populations has been calculated through *z*-score-standardising measurements of cranial dimensions, working out the diversity of each (standardised) dimension, and averaging this diversity across the dimensions (e.g., Betti et al., 2009; von Cramon-Taubadel & Lycett, 2008); the method of representing trendlines for different types of diversities in Figure 2 was inspired by that approach.

## Bayesian information criterion and the origin of expansion

The way of estimating origins through the BIC (Betti et al., 2009; Manica et al., 2007) was used to determine origin areas (and therefore peak points). To find origin areas (and peak points), a previous study used 99 origins in Africa from Betti et al. (2013) and 32 coordinate pairs from von Cramon-Taubadel and Lycett (2008) (Cenac, 2022b). When defining origin areas, the present study used those 99 in Africa and, of the 32 from von Cramon-Taubadel and Lycett (2008), the present study used the coordinates which were not labelled as being inside Africa (26 coordinate pairs) (von Cramon-Taubadel & Lycett used four of the 32 as origins, plus two other coordinate pairs as origins). As before (Cenac, 2022b), the four-BIC level (e.g., Manica et al., 2007) was i) the way of seeing if the best quadratic model was a better fit than the best linear one, and ii) if the quadratic one is better, estimating an origin area employing quadratic models. For linear models involving certain diversities (autosomal SNP haplotype, X-chromosomal, Y-chromosomal, and cranial shape), BICs calculated in Cenac (2022b) were used (including BICs regarding autosomal SNP haplotype heterozygosity, which were calculated with, and without, San – Cenac, 2022b, found San had low autosomal SNP haplotype diversity for their distance from some of the locations in Africa, so, regarding autosomal SNP haplotype diversity, analysis in the present study occurred in the presence and absence of San).

To see if there is any point in adjusting cranial size dimorphism for absolute latitude (in the context of expansion), previous research using linear models has compared BICs using African origins (the 99 from Betti et al., 2013), to see if any BICs found when adjusting cranial size dimorphism are notably lower (using the four-BIC condition) than the lowest BIC when dimorphism is not adjusted (Cenac, 2022b); the present study did the same, but with mitochondrial diversity in the place of cranial size dimorphism, and minimum temperature instead of absolute latitude. This was not only done amongst linear models but also amongst quadratic ones, and the aforementioned 26 non-African coordinate pairs were used too.

## Results and discussion

For autosomal microsatellite heterozygosity, the best quadratic model fit diversity better than the best linear model, i.e., a non-linear trend was supported at a global level (Figure 2). This adds some weight to the quadratic trend found within sub-Saharan Africa in Cenac (2022b). Like with a linear model (Manica et al., 2007; Figure 1A), the quadratic model still suggested an African origin of the expansion (Figure 1B). In somewhat of a contrast to analyses in sub-Saharan Africa (Cenac, 2022b), the trendline (Figure 2) did not indicate an increase at shorter distances. Yet, few populations were at such distances (Table 1) like in Cenac (2022b).

Within sub-Saharan Africa, autosomal microsatellite heterozygosity has seemed to have a non-linear association with distance from southern Africa – a potential rise of diversity, then a fall (Cenac, 2022b). So, in the current study, there may be an expectation that the non-linear trend regarding autosomal microsatellite heterozygosity worldwide (Figure 2) could be because populations in Africa were present in the analysis. For male cranial diversity, Betti et al. (2009) found whether non-linearity (regarding cranial diversity for males) was apparent in the presence/absence of Patagonian populations; in the current study, for an idea of which continent (whether Africa or otherwise) may be contributing to the non-linearity regarding autosomal microsatellite diversity worldwide, analysis comparing linear and quadratic models (using Africa as the origin) was repeated a number of times – each time was in the absence of one of the continents,^1^ with a focus on whether quadratic models are better than linear models when using distances from the peak point represented in Figure 1B. The quadratic model was better in the absence of either Africa, Asia, Europe, or North America. On the other hand, without South America, the quadratic and linear models were equally good. And so, the non-linearity regarding autosomal microsatellite heterozygosity (Figure 2) might have been because of the presence of South American populations (in the context of the other populations). Future research could examine if this stands when using more populations^2^ for which Pemberton et al. (2013) presented autosomal microsatellite heterozygosities – perhaps the research could use the 239 in Figure 6A of Pemberton et al., with which Pemberton et al. observed that heterozygosity declined whilst distance from eastern Africa rose.^3^

A non-linear trend was supported regarding mitochondrial diversity (unadjusted for minimum temperature) (Figure 2). Unlike autosomal microsatellite heterozygosity, the trendline with mitochondrial diversity indicated that the decline of diversity becomes weaker as distance increases before there is an increase in diversity (Figure 2) (like Betti et al., 2009, with male cranial form diversity prior to the removal of outliers from analysis). This appears to be comparable to the pattern in Figure 2C of Balloux et al. (2009). Through mitochondrial diversity, an origin area in Africa alone is apparent in analysis using linear models (Cenac, 2022b), and this also happened with quadratic models (Figure 1C). However, compared to that area found via linear models (Cenac, 2022b), the area obtained through quadratic models does appear smaller (Figure 1C). The non-linear trend does not indicate an initial increase of mitochondrial diversity (Figure 2). Amongst linear models, and amongst quadratic ones, BICs suggested that there is no reason in adjusting mitochondrial diversity for minimum temperature. Nonetheless, when mitochondrial diversity was adjusted for minimum temperature, support was not given for using a quadratic model (over a linear model). Therefore, the non-linearity in the relationship between mitochondrial diversity and distance from Africa (Figure 2) may not actually be a product of the expansion, but of climate.

Backing up linear declines with diversities in autosomal SNP haplotypes, the X-chromosome (Balloux et al., 2009), and cranial shape (von Cramon-Taubadel & Lycett, 2008), a linear model was the most appropriate for autosomal SNP haplotype diversity (regardless of whether San were included), X-chromosomal diversity, and cranial shape diversity (Figure 2). For Y-chromosomal diversity, a quadratic trend did not appear. Therefore, the inability of Y-chromosomal microsatellite heterozygosity to indicate the expansion (Cenac, 2022b) would not appear to be because of non-linearity.^4^

Previous research suggested that peak points (for various diversities and cranial size dimorphism) broadly converge on the south of Africa, and a centroid calculated from peak points was in the south, hence supporting the south being the most likely region of Africa in which the expansion originated (Cenac, 2022b). The peak point found with autosomal microsatellite heterozygosity seems more southerly when using a quadratic model compared to a linear one (Figure 1). When calculating the centroid, the peak point from Ramachandran et al. (2005) for autosomal microsatellite heterozygosity had been used (Cenac, 2022b). The peak point was found from linear models (Ramachandran et al., 2005). Given the present study (and reasoning in the note of Figure 1), it can be assumed that the peak point in Ramachandran et al. (2005) would be more southern had they used a quadratic model. Regarding mitochondrial diversity, the peak point from linear models (Cenac, 2022b) is not quite as southerly as the peak point from quadratic models (Figure 1C). Therefore, peak points do still appear to generally gather on southern Africa.

Any analysis in the present study (see above) featured less than half the number of populations Cenac (2022b) used in sub-Saharan African analyses, so the sub-Saharan African analyses in Cenac (2022b) featured more populations in Africa than the present study. Because an origin in Africa is suggested (Cenac, 2022b; Manica et al., 2007; Figure 1), having more populations in the continent should generally provide a better opportunity to find if a decline is absent at shorter distances from the origin. Nevertheless, the sub-Saharan African analyses had few populations located near the origin (Cenac, 2022b), and the same can be said in the present study with populations globally (Table 1). And so, it is unclear what pattern is present across shorter distances from the origin of the expansion.

## Conclusion

It seems like there are non-linear relationships between distance from Africa and both autosomal microsatellite diversity and mitochondrial diversity (Figure 2), however i) the non-linear trend with autosomal microsatellite heterozygosity may arise from the inclusion of South American populations, and ii) support with mitochondrial diversity disappeared when this diversity was adjusted for minimum temperature. No support for non-linearity was evident with autosomal SNP haplotype, X-chromosomal, and cranial shape diversities. Additionally, the lack of synergy between expansion from Africa and Y-chromosomal microsatellite heterozygosity (Cenac, 2022b) does not seem to be explainable by (quadratic) non-linearity. For autosomal microsatellite and mitochondrial diversities, quadratic models favoured Africa having been the start of the expansion. Therefore, non-linearity would not appear to influence where the origin seems to be (cf. Liu et al., 2006) at the continental level. However, depending on whether linear or non-linear models are used, there can be a notable difference in which area of Africa is suggested as the origin (Figure 1A and 1B).

## Footnotes

1 From coordinates (Balloux et al., 2009) and maps (Bartholomew illustrated reference atlas of the world, 1985; López Herráez et al., 2009), it was already known which continents the population coordinates were in when calculating geographical distances in Cenac (2022b).

2 Cenac (2022b) and the present study used the Pemberton et al. (2013) heterozygosities; regarding linearity/non-linearity, Cenac (2022b) used 104-106 populations in sub-Saharan Africa, whereas the present study used 50 worldwide.

3 When the autosomal microsatellite heterozygosity of the 239 populations is displayed on a graph with distance from Africa (heterozygosity on the *y*-axis, distance on the *x*-, and a linear trendline) (Pemberton et al., 2013), it does not seem probable that non-linearity is present outside of Africa; nonetheless, looking at Figure 6A in Pemberton et al. (2013), heterozygosity appears relatively more dispersed (around the linear trendline) in the Americas (given the colour-coding used in figures in Pemberton et al., 2013) – so there might be heteroscedasticity. Amongst Native Americans, gene diversity has a positive correlation with non-Native-American ancestry (such ancestry being African / East Asian / European) (Hunley et al., 2016). Therefore, also considering the variation in non-Native-American ancestry amongst Native Americans (Hunley et al., 2016), it could be that heterozygosity in the Americas may possibly be widely dispersed in Pemberton et al. (2013) as a result of variation in non-Native-American ancestry in the Americas.

4 It has been observed that Y-chromosomal microsatellite diversity in Balloux et al. (2009) would not seem to be at its numerical greatest for Africa (Cenac, 2022b). Studies have concerned Y-chromosomal diversity regarding Africans and others (Wilson Sayres et al., 2014) (also see Jorde et al., 2000). For example, in Hammer et al. (2001), haplotype heterozygosity was at its numerical highest with Central Asians, however, when pairwise haplotype differences were averaged, sub-Saharan Africans had the greatest diversity numerically.

